# Phasic alertness generates urgency and amplifies competition between evidence accumulators

**DOI:** 10.1101/2024.06.18.599522

**Authors:** Jeshua Tromp, Franz Wurm, Federica Lucchi, Roy de Kleijn, Sander Nieuwenhuis

## Abstract

Although phasic alertness generally benefits cognitive performance, it often increases the interference caused by distracting information, resulting in impaired decision-making and cognitive control. However, it is unclear why phasic alertness has these negative effects. Here, we present a novel, biologically-informed account, according to which phasic alertness generates an evidence-independent urgency signal. This urgency signal shortens overall response times, but also amplifies competition between evidence accumulators, thus slowing down decision-making and impairing cognitive control. The key assumptions of this account are supported with pupil measurements and electrophysiological data from human decision-makers performing an arrow flanker task. We also show that a computational model of the flanker task that incorporates time-varying urgency can reproduce the behavioral effects of phasic alertness, but only when the evidence accumulators compete with each other through lateral inhibition. Our results reveal a close interplay between dynamic changes in urgency, cognitive control and evidence accumulation.

## Introduction

The alerting system, one of the attention systems of the human brain, plays a crucial role in our ability to rapidly process and act upon abrupt changes in the environment ^1,2^. Auditory and visual alerting cues that directly precede a target stimulus often benefit performance by enhancing perception ^3–5^, boosting visual search ^6,7^ or speeding up behavioral responses ^8,9^, even when they provide no information about the response to be made to the target stimulus. These effects are commonly attributed to phasic alertness, a transient increase in arousal that enhances the readiness of the body and brain to respond to external stimuli.

Although the effects of phasic alertness on cognitive performance are generally beneficial, a marked exception concerns the deleterious effects of auditory and visual alerting cues on cognitive control. These effects occur in decision-making tasks in which individuals must ignore salient but misleading aspects of a stimulus in favour of less prominent but task-relevant information. In these tasks, phasic alertness *increases* the interference caused by the distracting stimulus elements, even though overall decisions are made more quickly. This counterintuitive pattern of results has been reported in numerous studies in recent decades ^10–20^. However, it remains unclear why alerting cues impair cognitive control in these contexts: previous theoretical attempts to explain the relevant observations ^11,16,21,22^ have met with conflicting findings ^17,22–25^, and little is known about the neuroscientific basis of the interaction between alerting and cognitive control.

Here we present a novel, biologically informed account of how phasic alertness impacts decision-making and cognitive control (Fig. 1). The account consists of three key assumptions, which we support with empirical evidence and computational modeling. First, we assumed that alerting cues would cause a transient increase in phasic arousal ^26^, driven by neuromodulatory nuclei of the ascending arousal system, including the locus coeruleus ^27,28^. We tested this assumption by measuring the effect of auditory alerting cues on the pupil dilation response (PDR), a well-validated correlate of phasic activity of these neuromodulatory nuclei ^29,30^.

**Figure 1.**
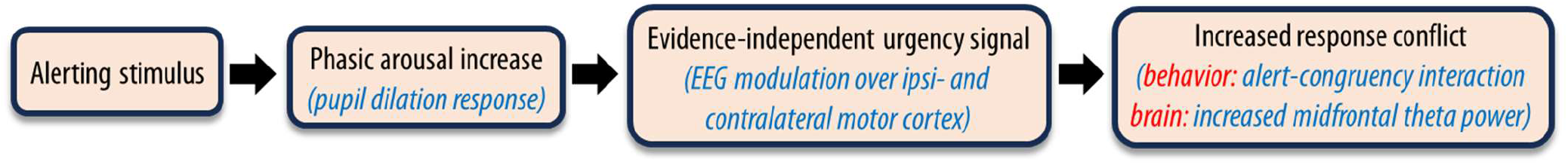
Hypothesized causal pathway (and operationalizations, in blue text) explaining the deleterious effect of phasic alertness on cognitive control. Black arrows depict the three key assumptions.

Second, we assumed that the phasic arousal response to alerting stimuli would produce an evidence-independent neural urgency signal that expedites the evolving decision process by driving evidence accumulators closer to a fixed decision threshold. If this assumption were correct, then the urgency signal caused by alerting stimuli would form an exogenous (“stimulus-triggered”) variant of the endogenous urgency signal that human subjects can invoke to speed up their decisions under time pressure ^31,32^. Several studies have found that decisions under time pressure are accompanied by larger PDRs ^31,33–35^. Furthermore, a pharmacological study in humans has revealed a causal effect of noradrenergic activity on (endogenous) decision urgency ^36^, providing further evidence for a link between neuromodulatory arousal systems and urgency. To examine this link in the context of phasic alertness, we assessed the effects of alerting cues and phasic, pupil-linked arousal on an EEG signature of decision urgency in motor cortical activity.

Third, we assumed that decision urgency would increase response conflict. By driving accumulators closer to a fixed decision threshold, urgency signals speed up decisions. However, if the accumulators representing the potential outcomes of a decision process compete with each other through lateral inhibition ^37^, then urgency might also exacerbate the interference from distracting information, thus increasing response conflict and impairing cognitive control. We tested this assumption by examining the effects of alerting cues and decision urgency on behavioral and neural manifestations of response conflict: the size of the flanker congruency effect, as indexed by response times and accuracy; and the amplitude of midfrontal theta-band activity around the time of the response ^38,39^.

The behavioural, pupillometric and electrophysiological findings reported below provide strong, convergent evidence for our account of how phasic alertness impacts decision-making and cognitive control. We also show that a drift diffusion model of the flanker task with incorporated urgency can reproduce the behavioral effects of alerting, but only when the evidence accumulators compete with each other through lateral inhibition.

### Auditory alerting cues speed up responding and increase flanker interference

Participants (N=60) performed 640 trials of an arrow flanker task (Fig. 2a), in which they were to identify the direction of the middle arrow (target) using their left or right index finger, as quickly and accurately as possible. On congruent trials (50%), the four flanker arrows pointed in the same direction as the target arrow; on incongruent trials (50%) they pointed in the opposite direction. In addition, on half of the trials, we presented a brief, uninformative auditory alerting cue starting 500 ms before stimulus onset. We chose this cue-target interval because it is the most-used interval in alerting-control studies, and because it allowed us to distinguish the pupil and EEG responses associated with the alerting cue and flanker stimulus. The mixed model equations we used to analyze the data are described under *Methods* and cited below.

**Figure 2.**
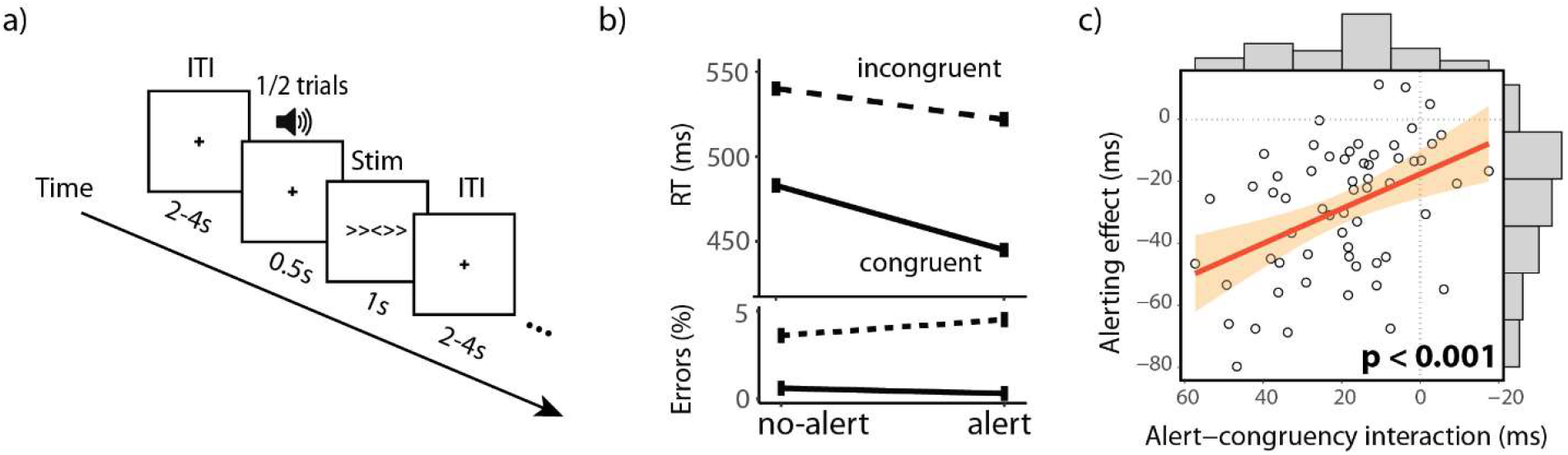
Task and behavioral results. a) Example of an incongruent trial in the arrow flanker task. The cue was a 150-ms, 800-Hz, 77-dB tone. b) Median reaction times and error rates as a function of congruency and alert condition. c) Scatterplot of alerting effect and alert-congruency interaction per participant, with distributions on top and right showing the interindividual variability in these effects. P-value taken from mixed model regression described in the main text. ITI = intertrial interval.

Median correct response times (RTs; Fig. 2b; Table 1) showed the expected effects of alerting, congruency and their interaction: RTs were slower on incongruent trials than on congruent trials (Equation 1a: β = -0.57, 95% CI [-0.59, -0.55], t(36813) = -48.60, p < .001), and faster on alert trials than on no-alert trials (β = -0.18, 95% CI [-0.20, -0.16], t(36813) = -15.26, p < .001). Importantly, we also replicated the typical alert-congruency interaction effect: the congruency effect was significantly larger on alert compared to no-alert trials (β = -0.20, 95% CI [-0.23, -0.16], t(36813) = -11.87, p < .001). This effect of alerting on the congruency effect was especially large on trials with a relatively small baseline pupil (Equation 2, three-way interaction: β = 0.03, 95% CI [0.007, 0.06], t(16689) = 2.56, p < 0.05) which are presumably characterized by relatively low baseline arousal, thus offering more scope for a strong phasic arousal effect ^40^. Furthermore, at the participant level, we found a significant relationship between the magnitude of the alerting effect and the alert-congruency interaction (Fig. 2c; β = 0.56, 95% CI [-0.86, -0.26], t(58) = 3.73, p < .001), suggesting that these effects reflect a common underlying process.

**Table 1.** Median response times and error rates (SD)

|  | Alert |  | No-alert |  |
| --- | --- | --- | --- | --- |
|  | Cong | Incong | Cong | Incong |
| RT (ms) | 428<br>(83) | 507<br>(95) | 462<br>(96) | 521 (102) |
| Err (%) | 1.3<br>(0.1) | 6.1<br>(0.2) | 2.8<br>(0.2) | 6.3 (0.2) |

As expected, error rates (Fig. 2b; Table 1) were larger on incongruent trials than on congruent trials (Equation 1b: *β* = -0.85, 95% *CI* [-1.00, -0.70], *p* < .001). Overall, alerting did not lead increases in error rates (*β* = -0.03, 95% *CI* [-0.15, 0.09], *p* = 0.60). Importantly, the congruency effect was increased on alert trials (*β* = -0.75, 95% *CI* [-0.99, -0.51], *p* < .001), mirroring the RT findings.

### Alerting cues elicit a phasic arousal increase

To assess if alerting cues elicited a phasic arousal increase, we compared the average pupil waveforms obtained in the four task conditions using a sliding-window mixed-model approach with a window length of 100 ms (N=31; Fig. 3). A significant alerting effect on the PDR emerged around 400 ms after the alerting cue (and 100 ms before the onset of the flanker stimulus) and lasted for 2 seconds. A significant effect of congruency, consistent with previous studies ^41^, emerged much later in the trial, around 900 ms after the onset of the flanker stimulus. We used these waveforms to extract a pupillary single-trial measure of phasic arousal ^26^ for use in subsquent analyses. To rule out any contribution from the pupil response associated with the flanker stimulus, we took the average pupil size in the 200 ms after the onset of this stimulus ^42^.

**Figure 3.**
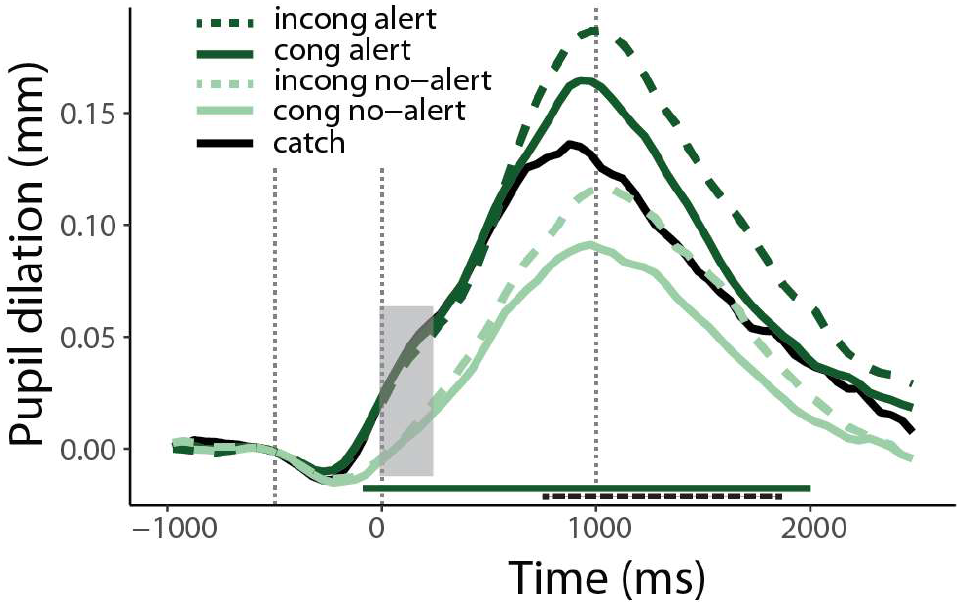
The alerting cue and congruency of the flankers impact pupil size at different times during the trial. Average pupil waveforms for each condition (correct trials only). The dark green horizontal bar indicates the time cluster that shows a significant main effect of alerting (p < 0.05). The black-striped horizontal bar indicates the time cluster showing a significant main effect of congruency (p < 0.05). The grey box indicates the time window chosen for further analyses. Vertical dotted lines indicate the onset of the alerting cues and the onset and offset of the flanker stimuli. incong = incongruent, cong = congruent; catch = the small proportion of catch trials, on which the alerting cue was not followed by a flanker stimulus.

### Alerting cues and pupil-linked arousal are associated with an EEG motor preparation signature of urgency

To test the second assumption of our theoretical account, we examined if alerting cues and the size of the PDR were associated with changes in decision urgency. Urgency modulations during decision-making can be measured in the motor cortex: neuronal activity in the premotor cortex and primary motor cortex reflects not only the accumulation of sensory evidence, but is also modulated by evidence-independent urgency before activity reaches a fixed firing-rate threshold ^43^. The same urgency signal also influences the vigor with which the ensuing response is made ^43,44^.

As a direct read-out of urgency modulations in the motor cortices ipsilateral and contralateral to the correct response hand, we computed the response-locked Laplacian-transformed EEG waveforms at electrodes C3 and C4 (Fig. 4a; N=53; ^45^). These waveforms showed an early activation of the ipsilateral (incorrect) motor cortex, reflecting the transient impact of incongruent flankers. This was followed by a rapid build-up of activity in the contralateral (correct) motor cortex, which typically peaks around EMG onset (not measured here) and is accompanied by inhibition of the ipsilateral motor cortex ^45^.

**Figure 4.**
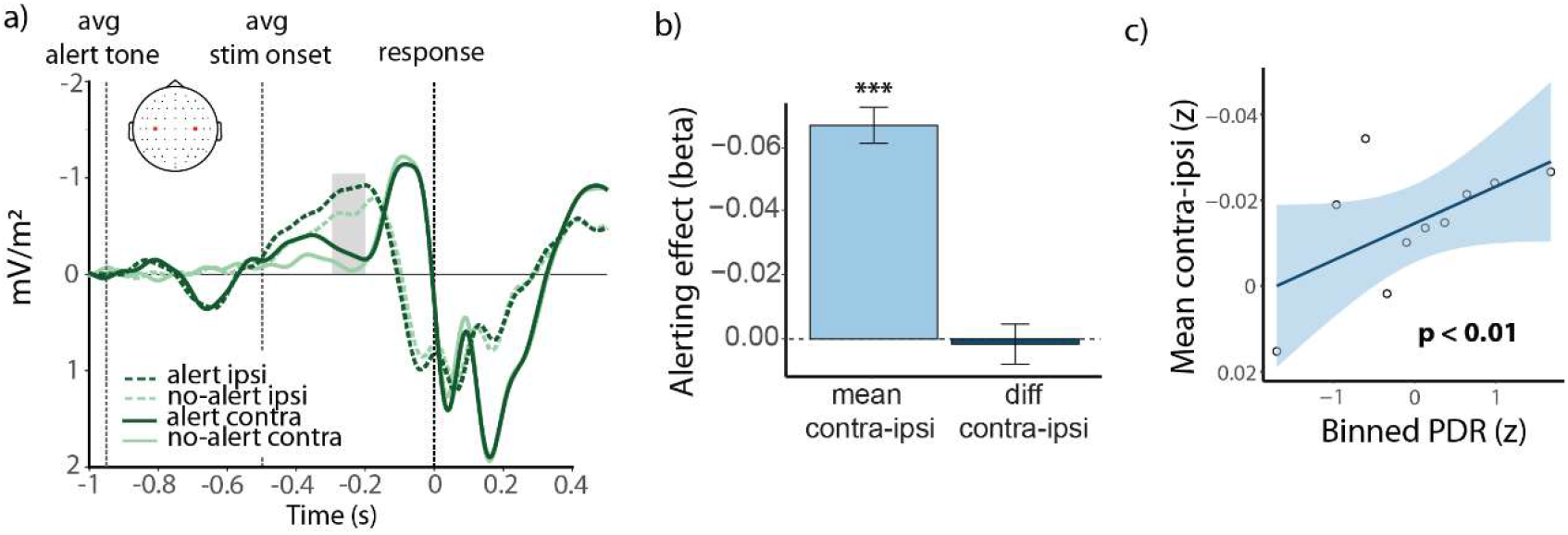
Alerting cues and pupil dilation response are associated with evidence-independent increase in motor cortex activity. a) Response-locked Laplacian-transformed EEG waveforms over the motor cortices ipsilateral and contralateral to the correct response hand (C3 and C4, red dots in scalp image). Waveforms are averaged across congruent and incongruent trials and are based on correct trials only. Note that stimulus-evoked potentials are smeared out in these response-locked waveforms due to variability in RTs. The grey box indicates the time window chosen for subsequent analyses. b) Statistical effect of the alerting cues on the mean of, and difference between, EEG activity over the contralateral and ipsilateral motor cortices. The beta values shown were derived from fits of Equations 3 and 4. c) Scatterplot showing the association between the size of the pupil dilation response and mean EEG activity over the contralateral and ipsilateral motor cortices. Shaded area indicates 95% confidence interval. ipsi = ipsilateral, contra = contralateral, PDR = pupil dilation response

Importantly, on alert trials the two motor cortices showed a similar-sized cue-evoked urgency modulation, which commenced around the (average) onset time of the flanker stimulus. To quantify the urgency modulations, we took the average amplitude of the waveforms in the interval from 300 to 200 ms before the button press. This interval prevented contamination of the measurements by stimulus-evoked potentials on trials with fast RTs. A statistical analysis confirmed the additive nature of the urgency modulation (Fig. 4b): the mean of contralateral and ipsilateral motor cortex activity increased after alerting cues (Equation 3: *β* = -0.06, 95% *CI* [-0.07, -0.04], *t*(28609) = -7.21, *p* < .001), while their difference did not (Equation 4: *β* =-0.004, 95% *CI* [-0.02, 0.01], *t*(28609) = -0.40, *p* = 0.69). Furthermore, focusing on participants who had both valid pupil and valid EEG data (N=29), we found that trials with a larger PDR were generally associated with larger mean motor cortex activity (Fig. 4c; Equation 5: *β* = -0.01, 95% *CI* [-0.02, -0.003], *t*(14866)= -1.15, *p* < 0.01), demonstrating a direct positive relationship between our measures of phasic arousal and decision urgency.

### EEG signature of urgency predicts speed of responding and flanker interference

Our account of how phasic alertness impacts decision-making and cognitive control assumes that urgency is the cause of both the speed-up of RTs and the increase in flanker interference. Therefore, our EEG signature of decision urgency should show this same relationship with RTs. Indeed, we found both a significant main effect of mean contra-ipsi on RT (Equation 6: *β* = 0.06, 95% *CI* [0.02, 0.09], *t*(28605) = 2.98, *p* < .01) and a significant interaction between mean contra-ipsi and congruency (Fig. 5a; *β* = 0.08, 95% *CI* [0.03, 0.13], *t*(28605) = 3.01, *p* < 0.01). Data generated by the best-fitting mixed model confirmed that RTs decreased with increasing bilateral motor cortex activity in both congruency conditions, but less so in the incongruent condition (Fig. 5b), suggesting that flanker interference increased with decision urgency.

**Figure 5.**
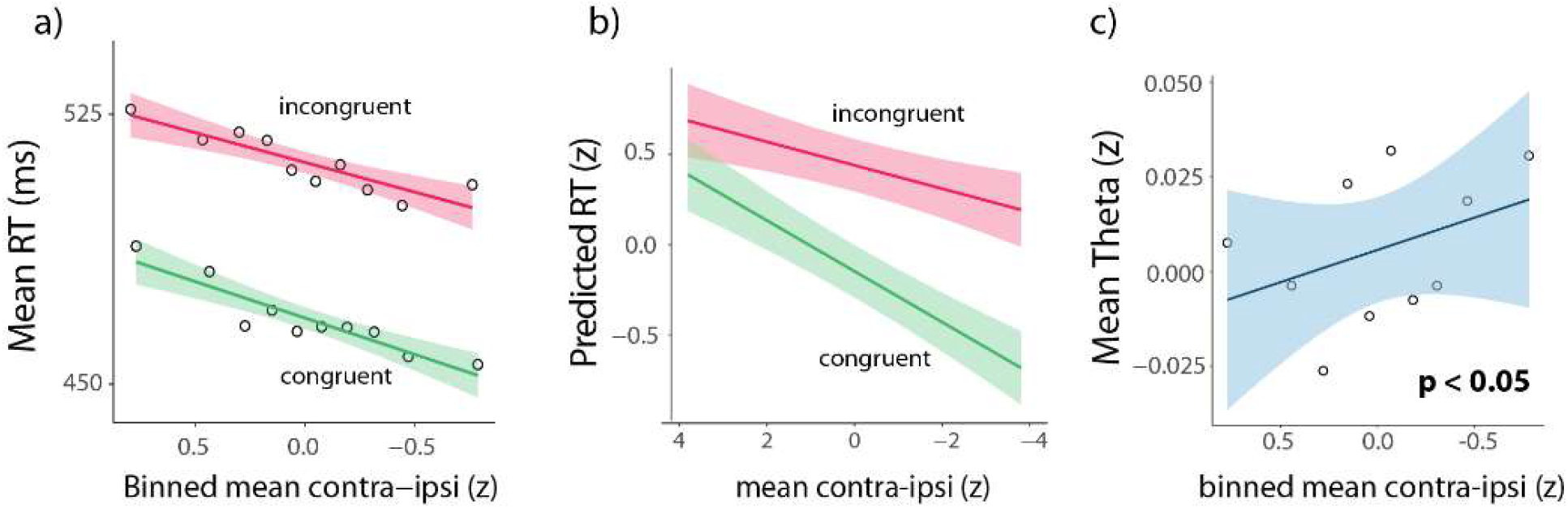
Higher bilateral motor cortex activity is associated with faster responses, larger congruency effects and higher midfrontal theta power. a) Scatterplot linking binned bilateral motor cortex activity to mean RT. b) Predicted RT for a range of mean contra-ipsi values, based on the fit of Equation 6, separately for congruent and incongruent trials. The range of the mean contra-ipsi values was chosen based on the minimum and maximum values in the empirical data. c) Scatterplot linking binned bilateral motor cortex activity to mean theta power. Note that negative x-axis values reflect higher motor cortex activity and thus we inverted x-axes in all panels. Shaded areas indicate 95% confidence interval.

### Alerting cues and EEG signature of urgency are associated with enhanced midfrontal conflict-related theta-band power

Next, we asked if the apparent increase in response conflict on alert trials (as reflected greater mean motor activity, and increased behavioural congruency effects) was accompanied by an increase in midfrontal theta power, an EEG index of response conflict ^38,39^. In line with previous research, we found an increase in midfrontal theta power around the time of the response (Fig. 6A; N=49). To examine the typical effect of response conflict on midfrontal theta power ^38,39^, we examined the incongruent – congruent difference waveform. The conflict effect revealed by this difference waveform peaked around the time of the response (Equation 7, main effect of congruency: *β* = -0.13, 95% *CI* [-0.16, -0.10], *t*(26759) = -7.98, *p* < .001) and was specific for the theta frequency range (Fig. 6B). Importantly, the conflict effect was stronger on alert trials than on no-alert trials (Fig. 6C; interaction: *β* = -0.10, 95% *CI* [-0.15, -0.05], *t*(26759) = -4.27, *p* < .001), supporting the assumption that alerting boosts the manifestation of response conflict in the brain ^46^.

**Figure 6.**
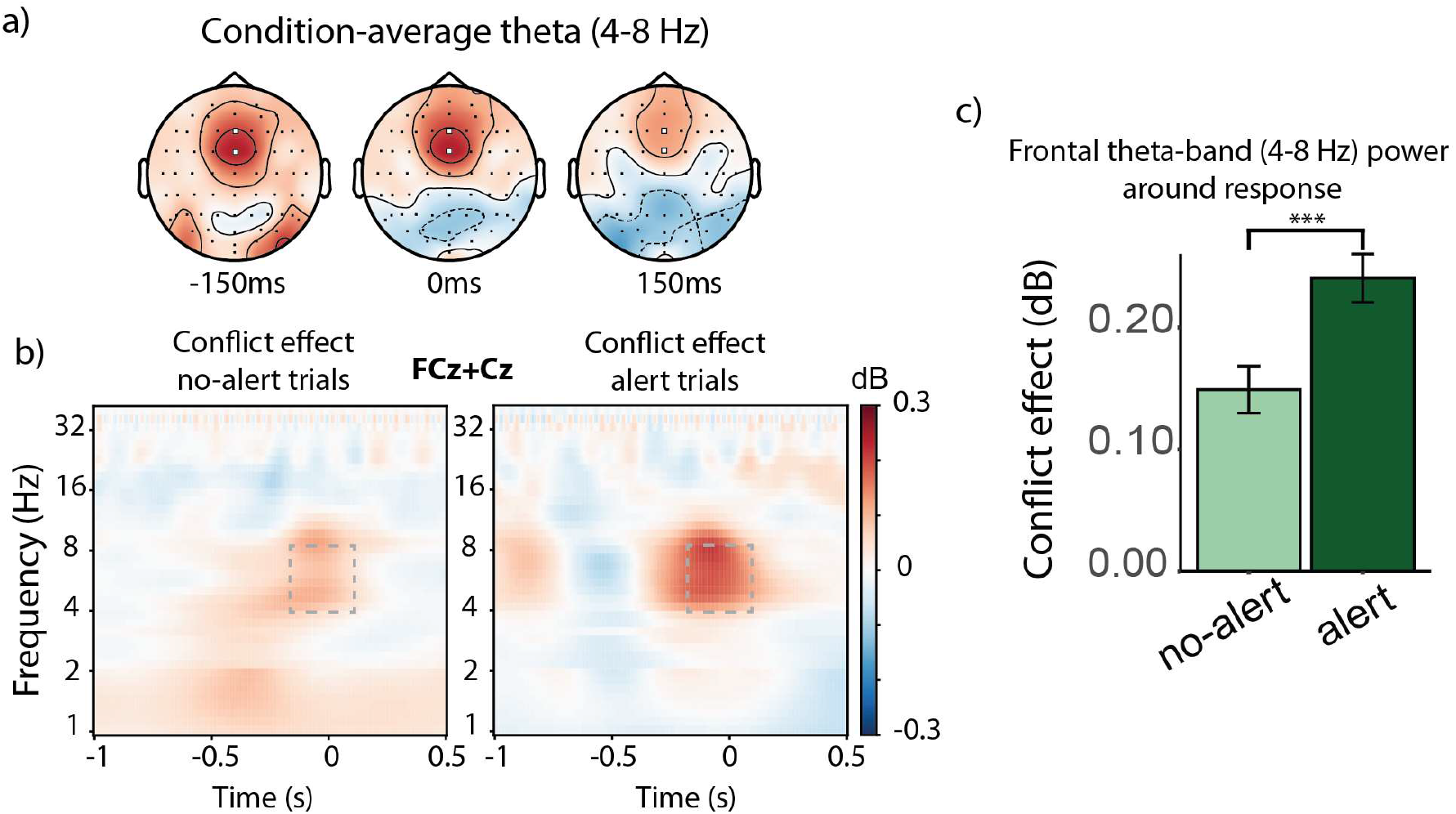
Midfrontal theta-band activity reveals increased response conflict in trials following an alerting cue. a) Response-locked scalp topographies showing scalp EEG activity in the theta band before, during and after the response. White points denote the midfrontal (FCz and Cz) electrodes. b) Time-frequency plot showing the response-locked conflict effect (incongruent – congruent) for no-alert trials (left) and alert trials (right). Dotted rectangles delineate the time-frequency range, based on van Driel et al. (2015), that was used for statistical analyses. c) Bar plot showing that the conflict effect is larger on alert trials than on no-alert trials. dB = decibel, Hz = Hertz.

Furthermore, corroborating our assumption that increased decision urgency leads to increased response conflict, we found a statistical relationship between bilateral motor activity and theta power (Fig. 5c; Equation 8: *β* = -0.03, 95% *CI* [-0.06, -0.004], *t*(26758) = -2.33, *p* < 0.05), but no relationship between the difference between contralateral and ipsilateral motor activity and theta power (Equation 9: *β* = 0.001, 95% *CI* [-0.03, 0.04], *t*(26758) = 0.49, *p* = 0.62).

### The effect of alerting cues on performance is partially mediated by pupil-linked arousal and EEG signature of urgency

Our theoretical account predicts that the effects of alerting cues on task performance are mediated, at least in part, by alerting effects on phasic arousal and decision urgency. To test this prediction, we conducted causal mixed-model mediation analyses, separately for PDR (Fig. 7a) and mean EEG over the two motor cortices (Fig. 7b). We found that there was indeed a significant average causal mediation effect (ACME) for both PDR (*ACME* = -0.004, *p* < 0.001) and mean contra-ipsi (*ACME* = -0.01, *p* < 0.001), supporting our hypothesized causal paths. However, the direct effect (DE) of alerting on RT was also strong in both analyses (PDR: *DE* = -0.18, *p* < 0.001; mean contra-ipsi: *DE* = -0.28, *p* < 0.001). We discuss the implications of this latter finding in the Discussion.

**Figure 7.**
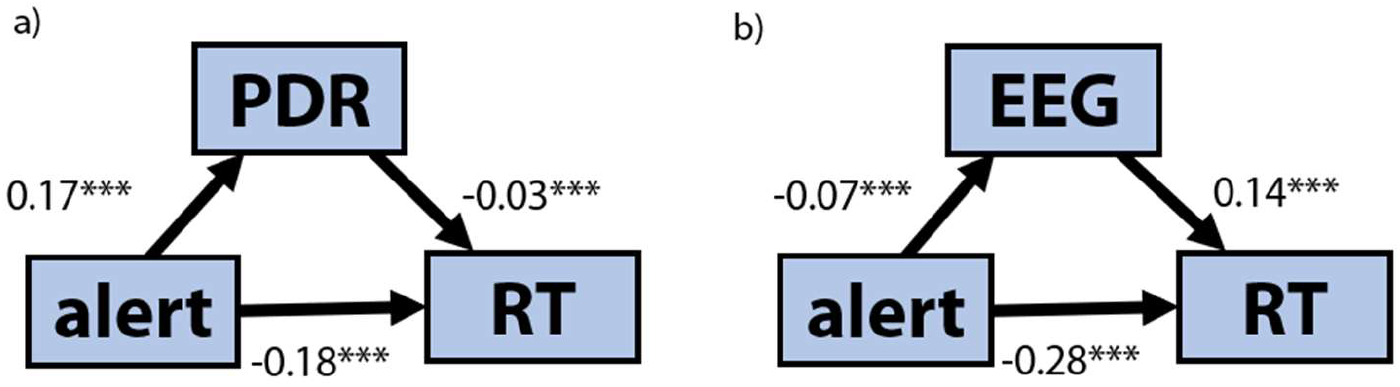
Alerting effects on response time are partially mediated by phasic arousal and mean activity over the contralateral and ipsilateral motor cortex. a) Causal mediation analysis diagram with a) PDR and b) mean contra-ipsi as mediators. PDR = pupil dilation response; EEG = mean contra-ipsi. *** denotes that the p-value is smaller than 0.001.

### Drift-diffusion modelling corroborates urgency account

We used drift-diffusion modeling to corroborate our assumption that injection of urgency into the decision process can reproduce the key qualitative alerting effects on flanker-task performance. In the regular drift diffusion model (DDM), drift rate is typically considered to remain constant over the course of a trial (aside from, in some model variants, intra-trial variability). To adapt the DDM for the flanker task, White et al. ^47^ proposed a shrinking spotlight (SSP) model, which assumes that attention in the flanker task is like a spotlight of which the width is gradually reduced over the course of a trial: it is diffuse at stimulus onset, allowing drift rate to be influenced by the flankers, but gradually narrows in on the central target, such that drift rate is increasingly dominated by the target. More formally, the overall decision evidence at any time point, drift rate *v*(t), is the sum of the perceptual strength of each stimulus item weighted by the amount of attention allocated to it, which is governed by the width of the spotlight at the beginning of the trial, and the rate at which the spotlight shrinks (see *Methods*). The SSP model has been shown to successfully account for various aspects of flanker-task performance ^47,48^.

The SSP model, like the regular DDM, has one accumulator which can drift toward either of two fixed decision boundaries, associated with the correct and incorrect response ^47^. To examine the effect of an evidence-independent urgency signal on behavorial performance, the DDM must be adapted so that it has two perfectly anti-correlated evidence accumulators that race toward the same decision threshold: if evidence for one accumulator increases, evidence for the other accumulator decreases by the same amount. In keeping with Weichart et al. ^49^, we refer to this variant of the SSP diffusion model with two accumulators with crossed inhibitory inputs as the feedforward inhibition (FFI) model (Fig. 8). To simulate the effect of alerting, the two accumulators were subjected to the same additive, time-varying urgency signal ^31^, thus constituting an additional input to the two evidence units compared to the no-alert condition. In line with the empirical urgency modulation in the EEG data (Fig. 4a), we assumed that urgency was already maximal at the start of the decision process (corresponding to *t*=0 in the model simulations), as characterized by a free (positive) intercept parameter. This moment corresponds to ∼150 ms after the average stimulus onset latency ^50^. Consistent with the transient nature of phasic alerting effects ^51^, the simulated urgency signal then diminished, as characterized by a free (negative) slope parameter.

**Figure 8.**
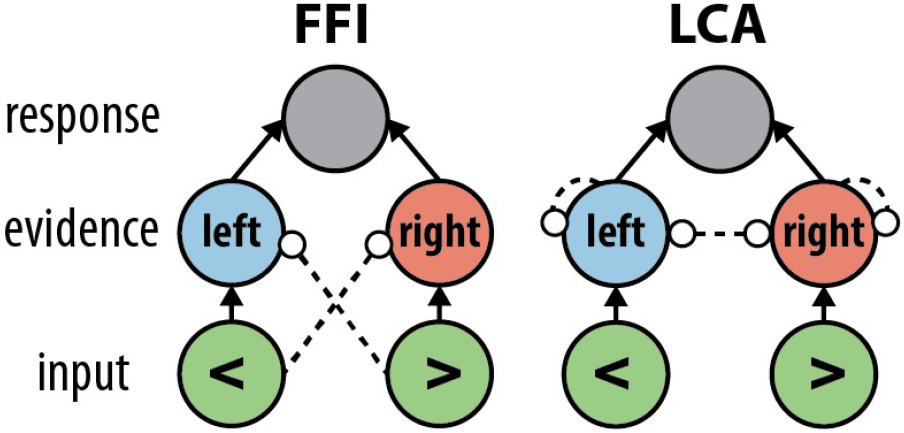
Schematics of the two models (Weichart et al., 2020). Solid arrows denote excitatory connections. Striped lines with circles denote inhibitory connections. FFI = feedforward inhibition, LCA = leaky-competing accumulator.

We also considered another variant of the SSP diffusion model, based on the leaky-competing accumulator (LCA) model of two-choice RT performance ^37,49^. Each accumulator in this LCA model variant passively leaks evidence over time, and is inhibited by the activation of the competing accumulator through lateral inhibition (i.e., instead of feedforward inhibition)(Fig. 8). Previous research found that the LCA model variant, with weakly anti-correlated evidence accumulators, provided a better account of behavioral and EEG flanker-task data than the FFI with strongly anti-correlated accumulators (which features no leakage) ^49^. To simulate the effects of alerting using this model, we injected the same additive, time-varying urgency signal as described above. We expected that if urgency causes the behavioral effects of alerting through amplification of the direct competition between the two evidence accumulators, then the LCA model but not the FFI model should be able to reproduce the empirical findings.

To simulate task performance, we first determined the set of parameter values for the FFI and LCA models that best fitted the behavioral data in the no-alert condition (median RT, SD of RT, accuracy; see *Methods* for fitting procedure), keeping the urgency parameters at zero. We then fixed these parameter values (Supplementary Table 1) and conducted a second fitting procedure to find the urgency parameter values (intercept, slope) that led to the best fit of the behavioral data in the alert condition. Figure 9 shows the performance in the no-alert condition of the best-fitting FFI and LCA models. Both models were able to capture key features of the empirical data: the congruency effect in both RT and accuracy (Fig. 9a), a skewed RT distribution with a fat tail (although both simulated distributions have fatter tails; Fig. 9b), and a conditional-accuracy function that shows a rapid decrease in the proportion of errors with increasing RT on incongruent trials (Fig. 9c), in line with the notion of a shrinking spotlight.

**Figure 9.**
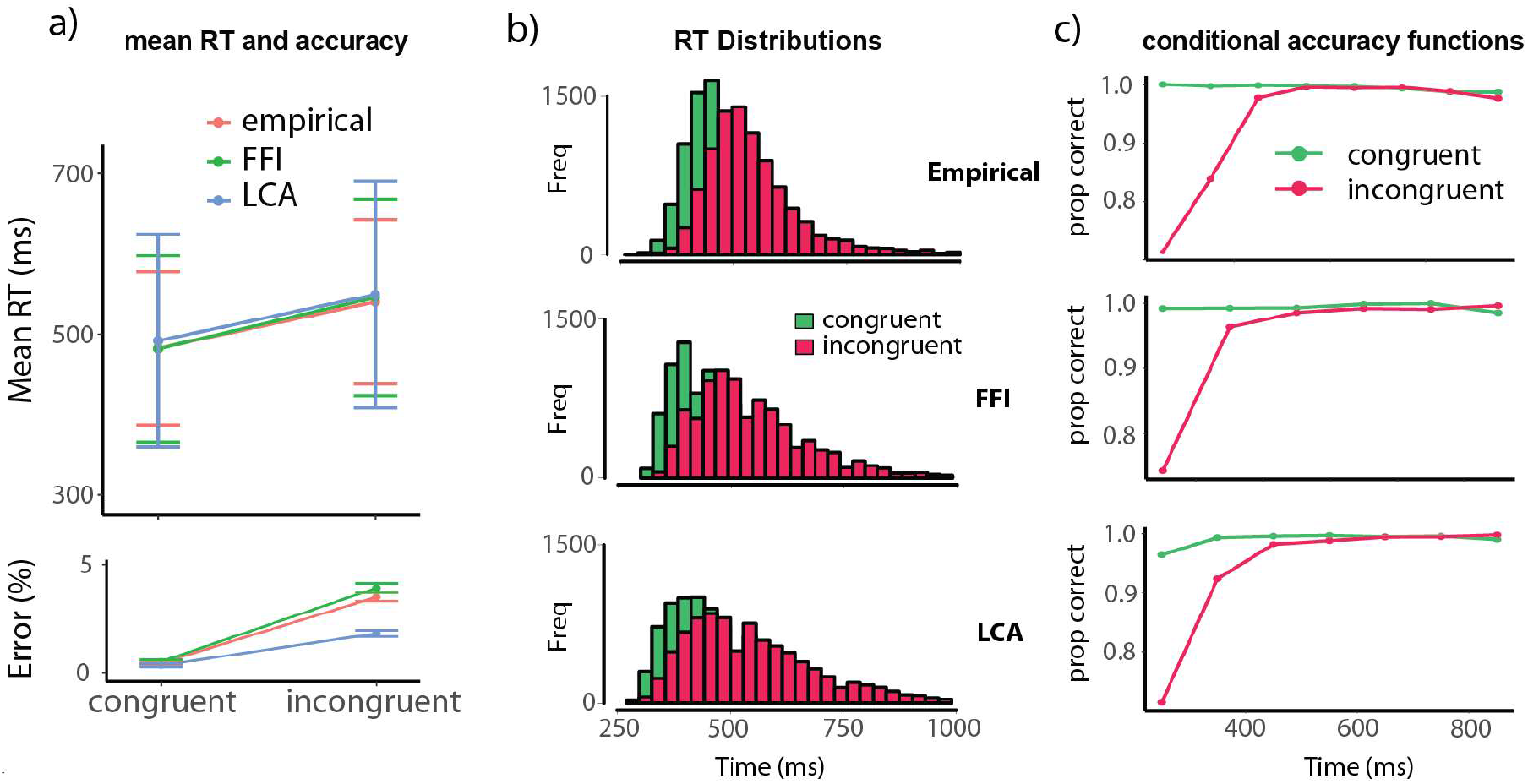
The FFI and LCA models provide good qualitative fits of empirical performance in the no-alert condition. a) Empirical and best-fitting simulated mean correct response times (top) and error rates (bottom) show typical congruency effects. Error bars depict standard deviations. b) Empirical and simulated correct response-time distributions. c) Empirical and simulated conditional accuracy functions show the typical increase in accuracy as a function of response time. FFI = feedforward inhibition, LCA = leaky-competing accumulator, freq = frequency.

In Figure 10, we compare the performance of the best-fitting FFI and LCA models with and without urgency. The best-fitting urgency parameter values (Supplementary Table 1) led both models to capture the general alerting effect on RT (Fig. 10a), but as expected only the LCA model could reproduce the alert-congruency interaction (Fig. 10b). This pattern of results was robust for a range of urgency parameter values (Supplementary Figure 3). The internal dynamics of the LCA model (Fig. 10c) confirm that urgency expedites overall RTs (i.e., the main effect of alerting) because it pushes the evidence accumulators closer to the decision threshold. The model also reproduces the main effect of congruency, because of the greater competition between the correct and incorrect accumulators on incongruent trials, especially at the start of the decision process (when the spotlight is still diffuse). This competition, through lateral inhibition, is especially pronounced in the alert condition because both accumulators receive an additional input through the urgency signal. This urgency modulation slowly dissipates through passive leakage and therefore lingers after the spotlight has converged on the target and the urgency signal has returned to zero (Fig. 10c), resulting in the larger congruency effect on simulated alert trials (i.e., the alert-congruency interaction effect). The FFI model also reproduces the main effects of congruency and alerting. However, because there is no lateral inhibition between the two evidence accumulators, the urgency inputs to the accumulators do not amplify their competition, and therefore do not result in an increased congruency effect on simulated alert trials.

**Figure 10.**
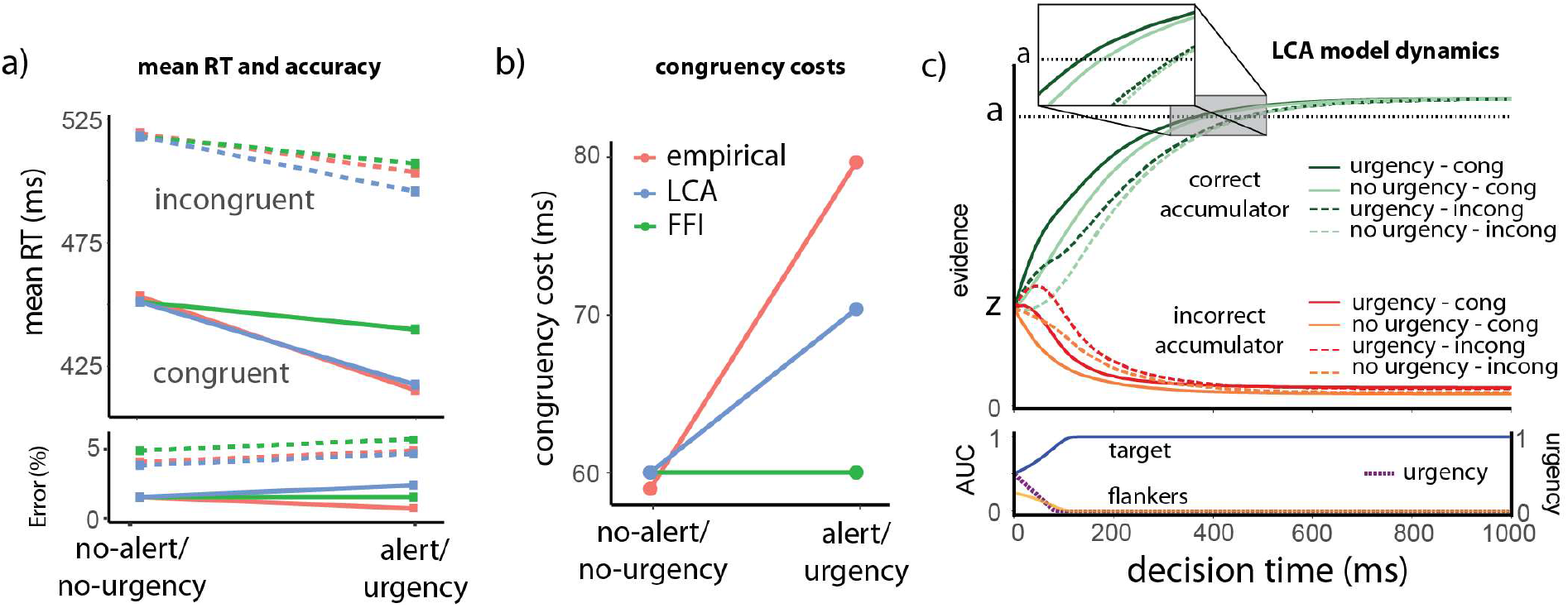
An LCA model with urgency can reproduce the pattern of empirical behavioral results. a) Empirical effects of alerting and best-fitting simulated effects of urgency on mean correct response times (top) and error rates (bottom). b) Only the LCA model with urgency is able to reproduce the alert-congruency interaction. c) Top: The average LCA model dynamics for both accumulators, separately for each condition. The inset zooms in on the area of the plot where the correct accumulators reach the threshold (between 300-500 ms). Bottom, left y-axis: the area under the curve (AUC), which corresponds to the amount of attention allocated to the target and flanker stimuli, which influences the drift rates of the correct and incorrect accumulators. Bottom, right y-axis: the best-fitting urgency signal. cong = congruent, incong = incongruent.

## Discussion

Although phasic alertness generally benefits cognitive performance, it increases the interference caused by distracting perceptual information, resulting in impaired cognitive control ^10–20^. The goal of the present research was to characterize the causal pathway underlying this subtle but ubiquitous interaction between alertness and control. Our results support the hypothesized causal pathway in Figure 1, which can be summarized as follows. First, alerting cues generate an evidence-independent urgency signal, as indicated by a phasic increase in pupil-linked arousal, a bilateral increase in motor cortex activity, and a general speed-up of responses. And second, this urgency signal amplifies competition between evidence accumulators and thus impairs cognitive control, as indicated by behavioral and neural markers of response conflict, and by simulations using an established computational model of the flanker task ^49^.

If an urgency signal drives evidence accumulators closer to their common decision threshold, this will not only expedite responses but may also increase the probability of an incorrect response ^31,32^. That is, distracting stimulus elements (e.g., flankers) and perceptual noise may lead an incorrect accumulator to cross the decision threshold before the correct accumulator. In line with this urgency effect on the speed-accuracy tradeoff, we found a minor but significant increase in error rates in the alert condition. Several perceptual decision-making studies have found a similar alerting effect on the speed-accuracy tradeoff, especially studies that used relatively simple speeded tasks (with very short RTs), in which the accumulators were presumably already close to the decision threshold in the no-alert condition ^8,12,52^. In contrast, in relatively complex tasks, in which the accumulators tend to start at a larger distance from the decision threshold, alerting cues push the accumulators closer to the decision threshold (speeding up responses), but not sufficiently close to risk more incorrect threshold crossings ^3^.

Although we propose a novel account of how phasic alertness impacts cognitive control, it is reminiscent of the facilitated response activation hypothesis ^20^, according to which “…, the presence of alerting signals is assumed to facilitate automatic stimulus-response translation processes for relevant and for irrelevant stimulus attributes, resulting in increased interference effects between simultaneously active response codes” ^53^; and which “… is based on dual-route frameworks of response preparation and proposes amplification of both direct response activation and indirect response selection processes” ^13^. That said, this hypothesis does not explictly refer to concepts such as arousal, urgency and evidence accumulation, and lacks the computational rigor and biological detail of the present work. In support of their response activation hypothesis, Fischer and colleagues reported that they only found an alert-congruency interaction in a word flanker task when the flanker words were drawn from the same set as the target words ^20^. This finding is not surprising from the perspective of our theoretical account, in which competition between evidence accumulators is an essential element. In terms of the LCA model in Figure 8: Alerting cues can only impair control if the distracting stimulus feature is processed by one of the competing pathways—either because that distracting stimulus feature also serves occasionally as a target (e.g., flanker arrows) or because it has an *ideomotor-compatible* effect on that pathway. For example, laterally presented incongruent Simon-task stimuli tend to drive the incorrect processing pathway because of the way in which the visuo-spatial-motor circuitry in the brain is hard-wired.

Our cue-target interval of 500 ms, although commonly used to study phasic alertness, was long enough to allow a bit of time for (voluntary) temporal preparation ^9,54^. This raises the question of whether some of the differences in results between the alert and no-alert conditions reflect in part an effect of temporal preparation instead of phasic arousal. Importantly, our mediation analyses revealed that the observed alerting effect on RT was significantly mediated by the PDR and by bilateral motor cortex activity, consistent with our proposal that the effects of alerting cues on task performance are mediated by phasic arousal and urgency. However, the mediation was partial, not full; both mediation analyses also revealed a significant direct effect of alerting on RT. Although we discouraged voluntary temporal preparation, using catch trials to weaken the temporal contingency between alerting cues and flanker stimuli, we cannot exclude the possibility that these direct effects were caused by temporal preparation. Indeed, preliminary evidence suggests that temporal preparation has similar effects on overall response speed and cognitive control as phasic alertness ^55,56^. Furthermore, recent evidence suggests that temporal preparation is accompanied by a gradual increase in pupil size and urgency ^57^. This raises the intriguing possibility that alerting cues can cause not only an exogenous, rapid and transient (i.e., phasic) increase in arousal and urgency, but also an endogenous, well-timed, relatively slow increase in arousal and urgency over the course of a predictable cue-target interval.

Phasic changes in pupil-linked arousal are known to be regulated by several neuromodulatory systems– in particular the noradrenergic system ^30,58^–that are thought to control global neural gain ^59,60^. This aligns with our urgency account: First, computational modeling studies have identified neural gain modulation, which regulates the responsivity of excitatory and inhibitory neurons, as a plausible mechanism for generating urgency in the brain ^31,61,62^. And second, although cortico-basal ganglia pathways are known to play an important role in the control of urgency ^32^, neural urgency signals can also be observed in choice-irrelevant sensorimotor regions ^63^, consistent with a role for the diffusely-projecting noradrenergic system in regulating urgency.

The largest challenge for any account of the alert-congruency interaction is that this interaction is not found for all conflict tasks. It is found in variants of the arrow flanker task ^12,22^, the Simon task ^13,14^, global/local task ^64^ and spatial Stroop task (i.e., classify the spatial meaning of a stimulus presented at an irrelevant position; ^19^. By far the most prominent example of a task that consistently fails to show the alert-congruency interaction is the manual color-word Stroop task ^16,24^. Darryl Schneider and other researchers have conducted many experiments to try to explain this dissociation. Most of the evidence suggests that the interaction only occurs in tasks requiring spatial attention and spatial information processing, and therefore not in tasks such as the Stroop task in which the relevant and irrelevant stimulus dimension are spatially integrated ^19,24^. However, why this might be the case is still poorly understood. Weinbach and Henik ^16^ proposed the attractive hypothesis that alerting causes a more diffuse focus of attention, resulting in more processing of spatially separate distracting information (e.g., flankers), and a corresponding increase in interference. However, this hypothesis, and two closely related accounts involving spatial attention ^17,21^ have not been supported in recent tests ^17,23,25^.

Alternatively, the distracting stimulus features in spatial conflict tasks (e.g., arrow flankers or the stimulus location in the Simon task and spatial Stroop task) tend to have directional associations (e.g., left/right or up/down) that can be mapped onto the spatial codes used in responding ^18,20,65^. Perhaps this is a necessary requirement for finding an alert-congruency effect. That is, an alerting cue may not boost activation along the incorrect pathway when the distracting stimulus feature (e.g., a color word) does not have directional associations with the response set (e.g., left and right response keys), as is the case in the manual color-word Stroop task ^24^.

However, the color-word Stroop task differs from spatial conflict tasks in other ways, which have not been considered yet in the context of interactions between alerting and cognitive control. For example, an analysis of the shape of RT distributions suggests that competition between evidence accumulators is fundamentally different in nature between the color-word Stroop task and spatial conflict tasks ^66^. Furthermore, the color-word Stroop task does not only elicit response conflict but also task conflict (i.e., reading the word versus naming the color; ^67^ which may interact with arousal and neural gain ^68^.

Moreover, we speculate that the neural network motif responsible for lateral inhibition (e.g., two excitatory populations of neurons representing evidence accumulators that mutually inhibit each other through a common population of inhibitory interneurons) may be more prominent in the spatial domain, given the topographical (i.e., spatial) organization of large parts of the cortex. Therefore, more work is needed to determine whether our urgency account offers an explanation for the discrepant results between conflict tasks. In any case, our urgency account predicts that compared to spatial conflict tasks the manual color-word Stroop task should be associated with strongly reduced urgency (at least in the incorrect pathway) and/or reduced competition between evidence accumulators, as evidenced by physiological measures.

There are several other outstanding issues. First, in the current work we focused on relatively small, stimulus-evoked fluctuations in alertness. An interesting question is how cognitive control is affected by intrinsic, relatively large and slow fluctuations in alertness during the waking state ^69^. Second, it is unclear how the conclusion that high alertness results in increased interference from distracting information can be reconciled with studies suggesting that phasic arousal tends to bias attention toward high-priority (e.g., perceptually salient or goal-relevant) stimuli ^70^. A potential explanation may lie in the knowledge that early in a flanker trial the bottom-up salience of the task-irrelevant information (i.e., the flankers) tends to dominate the competition between accumulators ^47,71^. If phasic arousal biases attention toward this task-irrelevant but highly salient information, this will result in larger flanker interference effects ^72^.

Finally, we used the SSP-LCA model to confirm the computational plausibility of our hypothesis that alerting effects on cognitive control tasks reflect an interaction between a shrinking attentional spotlight, a dynamic urgency signal and lateral inhibition between evidence accumulators. As in the work by Weichart and colleagues ^49^, this single-boundary dual-accumulator model with weakly anti-correlated evidence accumulators outperformed the FFI variant of the SSP model with strongly anti-correlated accumulators. However, it also has some limitations: It cannot accurately explain the presence of covert muscle activity of the incorrect response effector (i.e., partial errors; ^48^. And an SSP-LCA model variant in which the attentional spotlight width is based on within-trial response-conflict signals instead of simple passage of time has been found to account better for several aspects of flanker-task performance ^49^. Future studies should incorporate an urgency mechanism in this more complex SSP-LCA model variant and examine its potential to account for interactions between alerting and cognitive control.

## Aknowledgements

JT and SN were supported by the Netherlands Organization for Scientific Research (grant No. VI.C.181.032). The authors thank Peter Murphy for helpful comments on an earlier draft of the manuscript.

## Data availability

The datasets generated during and/or analyzed during the current study will be made available via the Open Science Framework (OSF) repository upon acceptance of this manuscript for publication. The Python scripts used for the model simulations, as well as the R script used for fitting the linear model are available on OSF: https://osf.io/vhq76/

## Competing interests

The authors declare no competing interests.

## Methods

### Participants

Sixty-four individuals (forty-one female), aged between 18 and 28 years, participated in the study in return for a monetary compensation of €14.25. They provided written informed consent, and the study was approved by the Psychology Research Ethics Committee of Leiden University (CEP code: S.T.-V3-2300). Four participants were excluded because they had an error rate of 25% or higher for one of the four trial types (see below), leaving sixty participants for further analyses.

### Task Design

Participants performed an arrow flanker task, in which the central arrow was flanked by two arrows on either side (Fig. 2a). The four flanking arrows pointed either in the same direction as the central target arrow (congruent trials) or in the opposite direction (incongruent trials). Each flanker stimulus remained visible for 1000 ms, irrespective of the participant’s response speed. We fixed the stimulus-viewing time to prevent a relationship between response time (RT) and the timing of visually-evoked pupil size changes. In half of the trials, an alerting tone (150 ms, 800 Hz, 77 dB) was played 500 ms before the stimulus (alert trials); in the other half, there was no tone (no-alert trials). To minimize previous-trial effects on pupil size, we let the intertrial interval (ITI) vary randomly between 2000 ms and 4000 ms.

The four trial types (congruent/alert, congruent/no-alert, incongruent/alert, and incongruent/no-alert) occurred equally often and were presented randomly. For each participant, there were ten blocks with 72 trials each. Eight were catch trials, in which an alerting tone was presented but no flanker stimulus. These catch trials were included to lower the temporal contingency between the alerting cue and the flanker stimulus, and thus discourage active temporal preparation during the cue-target interval (see *Discussion*).

### Procedure

We instructed participants to respond quickly and accurately to the direction of the central target arrow. Those who achieved an average RT of 400 ms or faster, with at least 95% accuracy, received an additional €4 compensation. Using a QWERTY keyboard, participants responded by pressing the “A” key with their left index finger for left-pointing arrows and the numeric pad’s “6” key with their right index finger for right-pointing arrows. We informed participants about the alerting cues but stressed that these were irrelevant to the task. Furthermore, we instructed participants to fixate their gaze on the central fixation cross presented between flanker stimuli.

Before the main experiment, participants completed a practice block of 16 trials and received feedback on their performance. Participants who did not achieve a 90% accuracy rate repeated the practice block until they did reach that accuracy. Participants were shown their average accuracy and RT after each block in the main experiment.

### Pupillometry

To minimize luminance-related pupil-size changes, we presented stimuli and background using isoluminant Teufel colors. The task featured a slate-blue background (RGB 166, 160, 198), salmon-colored arrows, and a salmon-colored central fixation cross (RGB 217, 152, 158). Using a Tobii-Pro eye tracker, we recorded pupil diameter under ambient light below 7.2 cd/m^2^ at a 40-Hz sampling rate.

Before the main experiment began, we calibrated the eye tracker. During the calibration and the main experiment, participants rested their heads on a chin rest placed 75 cm from the screen.

Pupil data from thirty-one participants were preprocessed using the *gazer* package in R. First, we identified eye blinks, which resulted in missing data during the blink and unreliable data shortly before and after the blink. For this reason, we also marked the 50 ms before and after the blink as missing data ^73^. The missing data were filled using linear interpolation, and the resulting data were smoothed with a moving-average filter with a width of 10 samples. All trials with over 50% missing samples were discarded, which amounted to 5.7% of the trials. Artifacts were identified based on the *max_pupil_dilation* function of the *gazer* package, which implements the outlier detection formula described by Kret and Shak-Shie ^74^. This led to the exclusion of an additional 0.4% of the trials. Lastly, we removed trials where the baseline pupil size deviated from the mean by more than three standard deviations ^73^, which led to the removal of 4.2% of the trials.

For the timepoint mixed-model regression analyses, we used a sliding window with a length of 100 ms to quantify the effect of trial type on the development of pupil size after the presentation of the alerting stimulus. These pupil time series were *z*-scored before they were entered into a mixed-model regression analysis—one for each window—with pupil size as the dependent variable, alerting and congruency as independent variables and subject as a random intercept to account for between-subject variability.

Informed by these analyses, we took the mean baseline-corrected pupil size in the interval between 0-200 ms after flanker stimulus onset as our pupil dilation response (PDR) measure for the trial-wise mixed model analyses. During this interval, pupil size showed a significant effect of alerting but not congruency. To baseline-correct the data, we used the interval from 600-500 ms before stimulus onset (i.e., the 100 ms before the potential presentation of the alerting stimulus).

Importantly, we found substantial time-on-task effects on the PDR within blocks but not over the whole experiment (see Supplementary Fig. 1). Because the PDR was substantially larger at the start of each block, we excluded the first five trials of each block before fitting the trial-wise mixed models described below.

### EEG data collection and preprocessing

Continuous EEG data were acquired using an ActiveTwo system (http://biosemi.com) from 64 scalp electrodes, configured to the standard 10/20 setup, and digitized at 512 Hz. Eye movements were recorded using two electrodes positioned above and below the left eye and two electrodes positioned at the outer canthus of each eye. Electrodes placed on the mastoids served as reference points.

We preprocessed the EEG data of sixty participants using a combination of manual and automatic preprocessing steps using the Python package MNE. First, we checked for electrode bridging. If bridging was present, we interpolated these electrodes. Next, we low-pass filtered the data at 0.5 Hz and cut the data into epochs around stimulus onset (-3500 ms – 2000 ms) and around the response (-3500 ms – 2000 ms). These wide time ranges prevented edge artifacts in the time-frequency decomposition described below. We then utilized an automated algorithm for unified rejection and repair of bad epochs called *autoreject*. First, we identified bad epochs using a first-pass *autoreject*, after which we conducted an ICA (using the Picard method) to filter out these bad epochs. This filter significantly increased the quality of the ICA components. After manual component rejection (mainly focused on eye-blink artifacts and other artifacts in the signal that we could clearly distinguish from brain activity), we re-ran the *autoreject* algorithm over all epochs, including trials excluded from the ICA. Finally, we visually inspected the EEG data. If key electrodes were still noisy or the *autoreject* algorithm rejected more than 30% of the epochs, we removed the participant from the analysis. This exclusion criteria led to removing seven of the sixty analyzed participants. Four additional participants were excluded from the midfrontal time-frequency analysis because of faulty/bridged frontal electrodes. A summary of each step of the preprocessing pipeline can be reviewed for each participant in the *osf* data storage (https://osf.io/an4sv/).

### EEG surface Laplacians over the motor areas

Next, we estimated the surface Laplacian (current source density) for response-locked epochs using a 10^th^-order Legendre polynomial. We set the regularization parameter (*λ*) to 10^5^ to optimize the balance between data representation fidelity and the smoothness of the estimated current source densities. This method accentuates local neural activities while minimizing the impact of volume conduction, thus enhancing the spatial resolution of the EEG data.

For the surface Laplacian analysis, we took the response-locked epochs and baselined these epochs by subtracting the mean of the baseline period 1700 ms – 1500 ms before the response from the entire signal. This time window ensured the baseline period was in the intertrial interval, regardless of response speed. We re-defined the signals of the C3 and C4 electrodes as ipsilateral and contralateral motor cortex signals, depending on which response was required. After visually inspecting the condition-average ipsilateral and contralateral waveforms, we identified the time window from -300 ms to -200 ms before the response as a suitable single-trial measure (see trial-wise mixed models below) for investigating urgency modulations of decision-related ipsilateral and contralateral motor cortex activity.

This time window prevented contamination of the EEG urgency measure by stimulus-evoked potentials on trials with fast RTs.

Note that in previous work we studied urgency by examining oscillatory power in the mu (8-14 Hz) frequency range, an alternative EEG measure of preparatory motor activity ^31^. However, in the current study, RTs were so fast that motor-related mu signals were contaminated by the influence of the stereotyped global decrease in 8-14 Hz power that tends to occur immediately after stimulus onset.

### EEG time-frequency analysis

For all time-frequency analyses, we decomposed the EEG time series into their time-frequency representations by convolving them with a set of Morlet wavelets and applying a fast Fourier transform to both the wavelets and the EEG data. The width of the Gaussian of the Morlet waves was set based on frequency: two cycles for frequencies < 2 Hz, four cycles for frequencies < 3 Hz, and six cycles for higher frequencies. These cycles yielded a good trade-off between temporal and frequency resolution.

For the response-locked time-frequency scalp plots (Fig. 6), we narrowed the frequency range to 4-8 Hz (i.e., theta) and split this range into ten log-spaced bins. We then applied a log-ratio baseline correction using a baseline period of 1700 ms – 1500 ms before the response, resulting in decibel-normalized power (dB). Next, we plotted the scalp topographies around the response (-150 ms, 0 ms, and 150 ms) and identified the mid-frontal electrodes (FCz and Cz) as the locus of increased theta power, in line with previous findings ^39^. Then, we ran a new time-frequency decomposition over the midfrontal electrodes with a broader frequency range. We increased the frequency from 1 to 40 Hz in 40 logarithmically spaced steps, with a subsequent log-ratio baseline correction. This time-frequency decomposition with a broad frequency range allowed us to test whether the conflict effect in the mid-frontal electrodes was specific to the theta band. For the trial-wise mixed-model analyses that include theta power, we selected the time window from -200 ms to 100 ms around the response (cf. ^39^. We calculated the mean dB for that time window for all frequencies between 4-8 Hz.

### Single-trial statistical analyses

We conducted the single-trial statistical analyses in R-Studio using the lme4 package for building and fitting linear mixed models. The complete code is available in the supplementary materials. For all mixed-model analyses, we filtered out the incorrect trials (except for Equation 1b). As independent variables, we took the mean of contralateral and ipsilateral motor activity (*mean contra-ipsi*), the difference between contralateral and ipsilateral motor activity (*diff contra-ipsi*), and response-locked theta power (*theta*), *z*-scored separately for each participant, as well as the sizes of the pupil dilation response (*PDR*) and baseline pupil (*BP*), *z*-scored separately for each participant and block. As dependent variables, we took reaction time (*RT*), *mean contra-ipsi, diff contra-ipsi*, and *theta*. These dependent variables were *z*-scored over all participants because the mixed models already took into account between-subject differences in the dependent variables; each model included a random intercept for each participant. To highlight the important results of these mixed models, we visualized them by dividing the independent variable into ten equally populated bins and showing the correlation with the dependent variable. Note, however, that the statistical tests were based on single-trial data, not binned averages. All mixed models are described using the R notation style.

First, we ran the following mixed model:

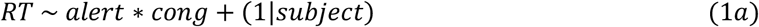

where *alert* denotes whether an alerting cue preceded the trial, and *cong* denotes whether the trial was congruent. The interaction term allowed us to test the key hypothesis that the congruency effect on RT (partly) depends on the alerting cue.

Second, we ran the following model:

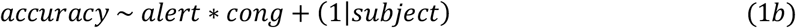

where *accuracy* is a binary variable denoting whether the response on a trial was correct. As the *accuracy* variable is binary, we ran a mixed model with a binomial distribution (in all the other models, we used the standard Gaussian distribution).

Next, we investigated pupil-related measures:

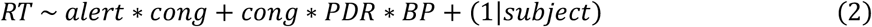

Followed by mixed models investigating the *mean contra-ipsi measure*:

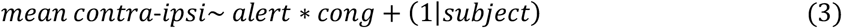

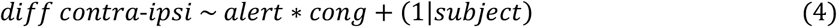

We predicted that alert trials would be characterized by a bilateral, not unilateral, increase in motor cortex activity. If that is the case, *mean contra-ipsi* should show an association with the alerting cue, but *diff contra-ipsi* should not.

To test our prediction that the bilateral increase in motor cortex activity depends on the size of the PDR, we ran the following mixed model:

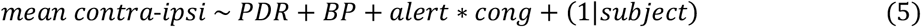

Next, we tested whether we could predict RT from *mean contra-ipsi*:

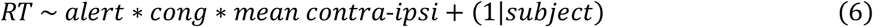

Then, we tested whether theta power at the time of the response was modulated by congruency, the alerting cue, and their interaction:

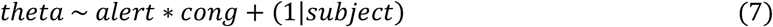

Subsequently, we tested the relationship between theta power and *mean contra-ipsi* versus *diff contra-ipsi*, with the prediction that theta should depend on the former but not the latter:

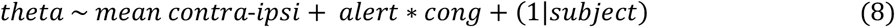

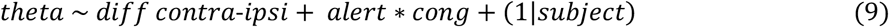

Lastly, we ran causal mixed-model mediation analyses ^75^ to assess whether *PDR* and *mean contra-ipsi* mediated the effects of the alerting cue on RT.

### Computational modeling

For the computational modeling, we took a previously successful sequential-sampling model of the flanker task as a starting point: the shrinking spotlight (SSP) model ^47,49^. The SSP model is a variant of the standard drift-diffusion model (DDM) for two-alternate decisions. The standard DDM model assumes that one accumulator accumulates noisy sensory input from a starting point *z* at a drift rate *v*. A decision is made when the accumulator reaches decision threshold *a* or *-a* (representing the two choices). All non-decision-related processing is captured in the non-decision time parameter *t*_*er*_.

The difference between the standard DDM and the SSP models we used is two-fold. First, the standard DDM assumes a constant drift rate over time, while the SSP includes a time-variant drift rate. The time-variant drift rate in the SSP is meant to implement the attentional spotlight hypothesis of the flanker interference effect, which assumes that attention gradually narrows in on the target during the trial. At stimulus onset, the spotlight is relatively wide, allowing the flankers to impact evidence accumulation; however, over the trial, attentional selection gradually reduces the spotlight width, leading the target arrow to dominate evidence accumulation. This hypothesis captures the commonly found below-chance drop in accuracy at fast RTs and other aspects of the conditional accuracy function ^71^. Second, the standard DDM has *one* accumulator with *two* decision boundaries, whereas the SSP model has *two* accumulators that race toward a single (common) decision threshold. Simulated trials on which the incorrect accumulator crossed the decision threshold first were considered error trials. This two-accumulator variant allowed us to add an urgency signal as evidence-independent input to both accumulators ^31^.

In the SSP model, the time-variant drift rate is governed by three parameters: perceptual input strength (*p)*, the width of the spotlight at the beginning of the trial (*sd*_*0*_), and the rate at which the spotlight shrinks (*r*_*d*_). The spotlight is a density function for a Gaussian distribution centered around 0, with standard deviation *sd*_*a*._ At each time point, the width of the spotlight is given by the following equation:

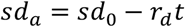

with *t* defining the time point of the trial, and *sd*_*a*_ defining the spotlight width at time point *t*. The resulting area of attention directed toward the target and the flankers is given by the following equation:

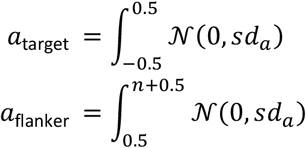

where *n* denotes the number of flankers, *a* denotes the area under the curve. Furthermore, we assume that each arrow (flanker or target) occupies one unit of space.

Finally, the drift rate is given by the following formulas, depending on congruency:

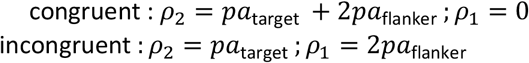

where *ρ*_2_ denotes the drift rate for the correct accumulator, *ρ*_l_ denotes the drift rate for the incorrect accumulator, and *p* denotes the perceptual input strength parameter.

We implemented two variations of the SSP model: the feedforward inhibition (FFI) variant and the leaky-competing accumulator (LCA) variant. These variants mainly differ in how the two competing accumulators relate. In the FFI variant, the accumulators are perfectly anti-correlated (cf. ^31^. This mimics the dynamics of the single-accumulator (standard) models, where a movement towards one decision threshold necessitates a movement away from the other. The following formula describes the accumulation of evidence in the FFI:

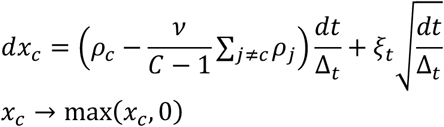

Evidence for each accumulator *c* is denoted *x*_*c*_ (with a lower bound of 0); *dx*_*c*_ denotes the change in evidence on timestep *t*; ρ_*c*_denotes the drift rate for accumulator *c*; ρ_*j*_denotes the drift rate for accumulator *j*, and noise is represented by ξ, a driftless Wiener process centered around 0.

In the LCA variant, the accumulators are not perfectly anti-correlated, but they inhibit each other to a certain extent through lateral inhibition, as given by parameter β. Additionally, in the LCA variant, the evidence in each accumulator passively decays throughout the decision process at a rate equal to leak parameter κ:

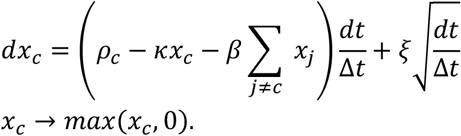

For further mathematical details of the SSP-FFI and SSP-LCA models, see Weichart et al. ^49^.

The approach we took was as follows. First, we fitted the two models to the empirical behavioral data in the no-alert condition. To determine the best-fitting set of model parameter values, we used a differential evolution algorithm to minimize a cost function that calculated, for each level of congruency, the difference between the empirical and simulated median RTs, between the empirical and simulated standard deviations (SD), and between empirical and simulated accuracy. The resulting differences were normalized by dividing by the empirical median RT, mean SD, and mean accuracy and then summed ^76^. We specified uninformative priors for each parameter (Supplementary Table 1). For each parameter set, we simulated 20.000 trials.

Our next step was to fit the two models to the empirical behavioral data in the alert condition. Therefore, we fixed all SSP parameters to the best-fitting values in the no-alert condition and added two free urgency parameters to simulate the effects of the alerting cue. The urgency signal was a time-variant function with a positive intercept and a slope, which added additional evidence-independent input to both accumulators evenly:

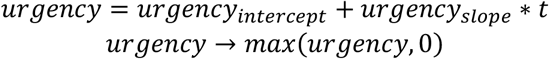

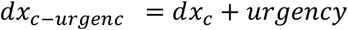

where *dx*_*c-urgenc*_ denotes the change in evidence for timestep *t*, including urgency. The urgency signal was constrained to be positive or zero. The model-fitting procedure was the same as for the no-alert condition. After acquiring the best-fitting urgency parameters for the FFI and LCA model variants, we conducted a grid search with 100.000 simulations per grid around the best-fitting urgency parameter combination to find out how robust the resulting urgency effect (average speed-up) and urgency-congruency effect (average increase in congruency effect due to the urgency) were (Supplementary Fig. 3).

## Notes

https://osf.io/vhq76/

### Competing Interest Statement

The authors have declared no competing interest.

