## Supplementary Files for "Phasic alertness generates urgency and amplifies competition between evidence accumulators"

**Supplementary figures**

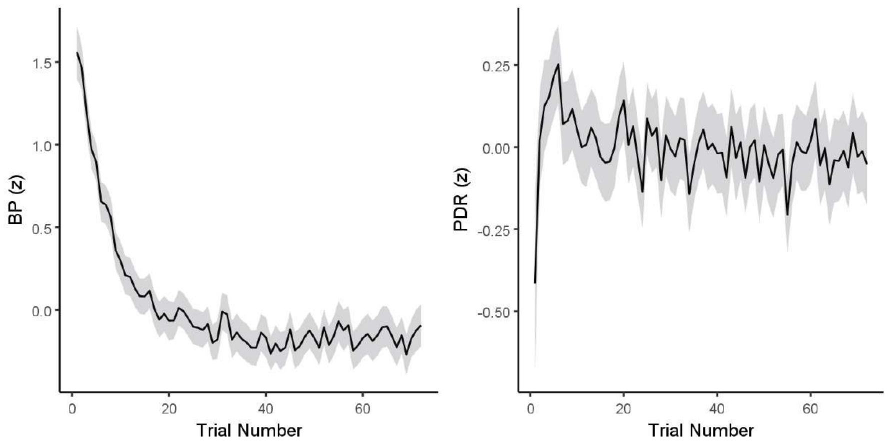


***Supplementary Figure 1. Pupillary measures deviate at the start of each block.*** *Left: baseline pupil (BP) as a function of trial number. Right: pupil-dilation response (PDR) as a function of trial number. The grey areas denote standard deviation. Both measures are z-scored per block per participant.*

.

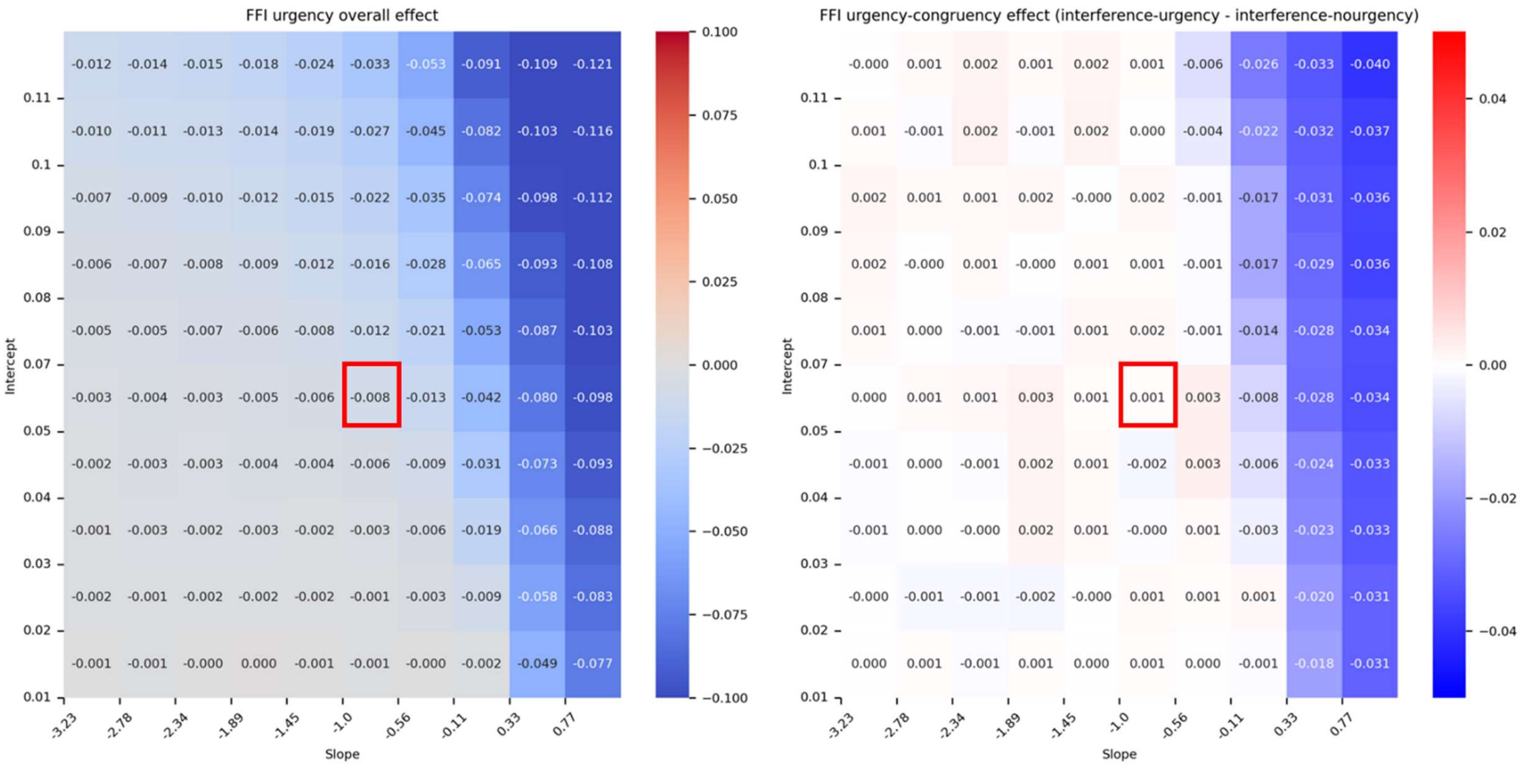


*Supplementary Figure 3. Grid search with 100.000 trial simulations per cell shows robustness of urgency-congruency findings in the FFI model. Left: overall urgency effect on RT averaged across congruent and incongruent trials. Right: urgency-congruency effect, calculated by subtracting the congruency effect on no-urgency trials from the congruency effect on urgency trials. A positive value thus means that there is increased interference in the urgency condition. The red squares show the best-fitting urgency parameters. See Methods for cost function and other aspects of fitting procedure.*

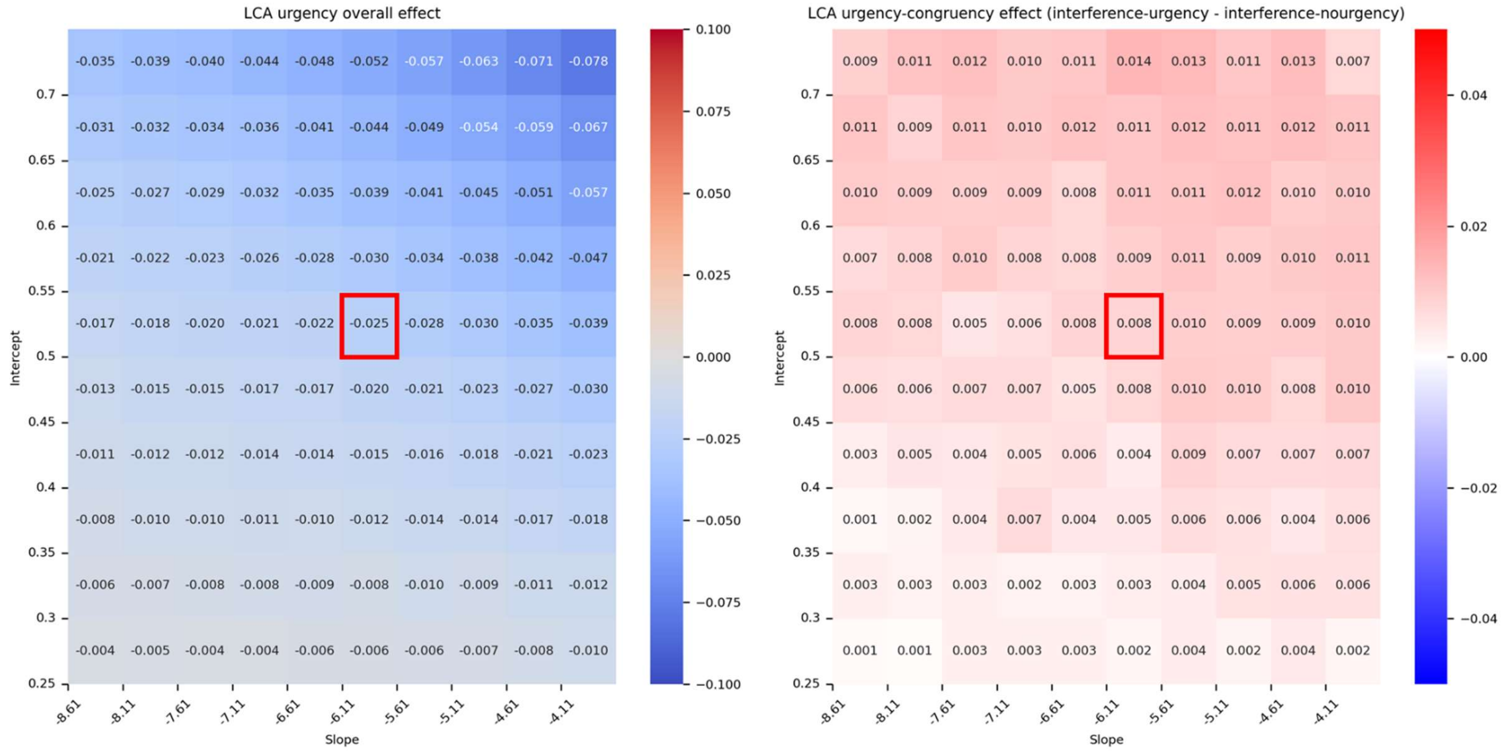


*Supplementary Figure 4. Grid search with 100.000 trial simulations per cell shows robustness of urgency-congruency findings in the LCA model. Left: overall urgency effect on RT averaged across congruent and incongruent trials. Right: urgency-congruency effect, calculated by subtracting the congruency effect on no-urgency trials from the congruency effect on urgency trials. A positive value thus means that there is increased interference in the urgency condition. The red squares show the best-fitting urgency parameters. See Methods for cost function and other aspects of fitting procedure.*

| Parameter | Description | Bounds | Best fit FFI | Best fit LCA |
|---|---|---|---|---|
| $r_d$ | rate of focus | (0,20) | 19.93 | 4.81 |
| $p$ | perceptual input strength | (0,20) | 1.88 | 6.22 |
| $sd_a$ | starting spotlight width | (0.5,5) | 1.72 | 0.73 |
| $a$ | decision threshold | (5,30) | 5.60 | 12.76 |
| $\xi$ | noise | (0.1,5) | 1.2 | 3.72 |
| $t_{er}$ | non-decision time | (0.1,0.25) | 0.25 | 0.24 |
| $\beta$ | lateral inhibition | (0.2,1) | - | 0.21 |
| $\kappa$ | leak | (0.2,1) | - | 0.63 |
| $urgency_{intercept}$ | urgency intercept | (0,5) | 0.05 | 0.5 |
| $urgency_{slope}$ | urgency slope | (-5,5) | -1.0 | -6.11 |

*Supplementary Table 1. Overview of parameters, bounds, and best fits for both FFI and LCA. The lateral inhibition and leak parameters are absent in the FFI models.*
